# StrainOptimizer empowers rational cell factory design through multi-scale metabolic models with expression and proteome constraints

**DOI:** 10.1101/2025.11.03.685948

**Authors:** Haoyu Wang, Mengyao Zhang, Chengyu Zhang, Siwei He, Wenbin Liao, Rongpeng Zhu, Yongjin J. Zhou, Hongzhong Lu

## Abstract

The rational design of microbial cell factories for high bioproduction remains a key challenge in metabolic engineering. While advanced modelling frameworks incorporating protein resource allocation, such as enzyme-constrained models (ecGEMs) and Expression and Thermodynamic Flux (ETFL), provide superior predictive power, their application is limited by a lack of user-friendly computational tools. Here, we present strainOptimizer, a comprehensive computational platform for rational strain design that systematically evaluates key resource allocation principles: the coupling of gene expression with metabolism, subcellular compartmentalization, and enzyme capacity limitations. Our benchmark analyses demonstrate that each principle offers distinct advantages: models coupling metabolism and expression (like ETFL) enable the identification of non-metabolic targets, organelle-level proteomic constraints improve precision for high–protein-cost products, and protein-usage-based objectives consistently outperformed traditional flux-based approaches. To demonstrate its practical utility, we applied strainOptimizer to an engineered sclareol-overproducing *Saccharomyces cerevisiae* strain. The platform identified novel targets, and experimental validation confirmed a 67% success rate, increasing the final sclareol titer by 14–26% and productivity by up to 45%. StrainOptimizer bridges the gap between resource allocation theory and applied engineering, providing a powerful, validated tool to accelerate the development of high-performance cell factories.

## Introduction

Biomanufacturing, which uses engineered microorganisms as cell factories, offers a sustainable and environmentally friendly route to produce a wide array of valuable chemicals (*1*). A central challenge in this field is the rational design of microbial strains to achieve cost-effective and high-yield production (*2*). While advances in synthetic biology have accelerated the genetic engineering cycle, the rational identification of effective modification targets remains a primary bottleneck due to the inherent complexity of cellular metabolism (*3*).

To address this challenge, systems biology and genome-scale metabolic modelling have provided powerful *in silico* frameworks for strain design. Over the past two decades, numerous classical computational algorithms—such as OptKnock (*4*), OptForce (*5*), FSEOF (*5*) and iBridge (*6*)—have been developed to guide metabolic engineering. Although valuable, these methods are built on classical genome-scale metabolic models (GEMs) that rely primarily on stoichiometric constraints. Consequently, their ability to predict cellular phenotypes is often limited, as they neglect critical biological limitations like finite enzyme capacity and proteome allocation (*7*).

More recently, the principle of protein resource allocation has been recognized as a fundamental concept linking metabolic fluxes to cellular physiology (*8*). Cells must optimally partition their finite proteome to meet both intrinsic functional demands (e.g., energy and precursor synthesis) (*9, 10*) and extrinsic physic constraint (e.g., cellular and organellar crowding) (*11, 12*). This has led to the development of enzyme-constrained GEMs (ecGEMs) (*11*), which incorporate proteomic constraints to more accurately capture complex phenotypes like metabolic overflow (*11*–*13*) and the hierarchical utilization of carbon sources (*14*).

The enhanced predictive power of ecGEMs has led to notable successes in metabolic engineering, including improved production of chemicals such as heme (*15*) and shikimate (*16*). However, these applications often rely on customized scripts, and the lack of accessible, user-friendly tools has hindered their widespread adoption. While several strain design algorithms for ecGEMs have been proposed, including ecFactory(*17*), ET-OptME(*18*) and so on(*18*–*20*), their development is often hampered by a lack of systematic pipeline evaluation and validation in practical experimental engineering contexts, making it difficult to convincingly demonstrate their advantages. For instance, our previously developed algorithm, ecFactory, provided valuable insights for constructing platform strains by analyzing 103 different chemicals (*17*). However, it was limited to a single model type, offered poor customizability, and lacked validation in a practical metabolic engineering case study. Recognizing these limitations, we were motivated to develop a next-generation computational platform that builds on the foundation of ecFactory to address these shortcomings.

More advanced multi-scale models that couple metabolic networks with gene transcription and translation—such as ME models (*21, 12, 22*) and ETFL models (*23, 13, 24*)—can simulate cellular states with even greater mechanistic accuracy. These models also include a broader range of genes, particularly non-metabolic genes, expanding the search space for novel engineering targets (*25*). However, despite this promise, their application in metabolic engineering remains rare. The primary bottleneck is a lack of adapted computational tools and strain design algorithms capable of managing their complexity and fully leveraging their unique predictive features. This motivated a systematic investigation into how their enhanced predictive power can be harnessed for *in silico* strain design.

Here, we present strainOptimizer, a comprehensive computational platform for cell factory design. First, we developed a modular platform and a high-quality benchmark dataset—comprising five diverse metabolic engineering case studies and a suite of evaluation metrics—to enable the comprehensive assessment and optimization of design algorithms. Using this platform, we systematically investigated how integrating protein allocation principles at three distinct levels—model types, additional constraints, and optimization objectives—impacts the performance of target prediction. Finally, to demonstrate its practical utility, we applied strainOptimizer to re-optimize an industrial-ready, high-yielding sclareol-producing *Saccharomyces cerevisiae* strain. By integrating strain-specific phenotype and transcriptomic data into our models, we predicted novel modification targets for experimental validation. Notably, 6 of 9 novel targets increased the final titer by 14% to 26%, with the highest productivity enhanced by 45%, demonstrating the platform’s effectiveness in a real-world engineering context. strainOptimizer bridges the gap between fundamental cellular principles and applied metabolic engineering, enabling the systematic and accurate prediction of genetic targets to enhance biomanufacturing performance.

## Results

### Framework of strainOptimizer

We developed strainOptimizer, a modular computational platform for systematic *in silico* strain design, extending the capabilities of our previous tool, ecFactory (*17*). To enhance its extensibility and accessibility for the research community, we migrated the entire codebase from MATLAB to Python, an open-source language with a more versatile and extensive scientific ecosystem (*26*). The platform was engineered with modular architecture, comprising five core components: Manipulation, Simulation, Score Rule, Analysis, and Strain Design Algorithms. This modularity provides users with significant flexibility, supporting both streamlined, one-click target predictions and the design of complex, customized computational workflows (Figure 1A).

**Figure 1.**
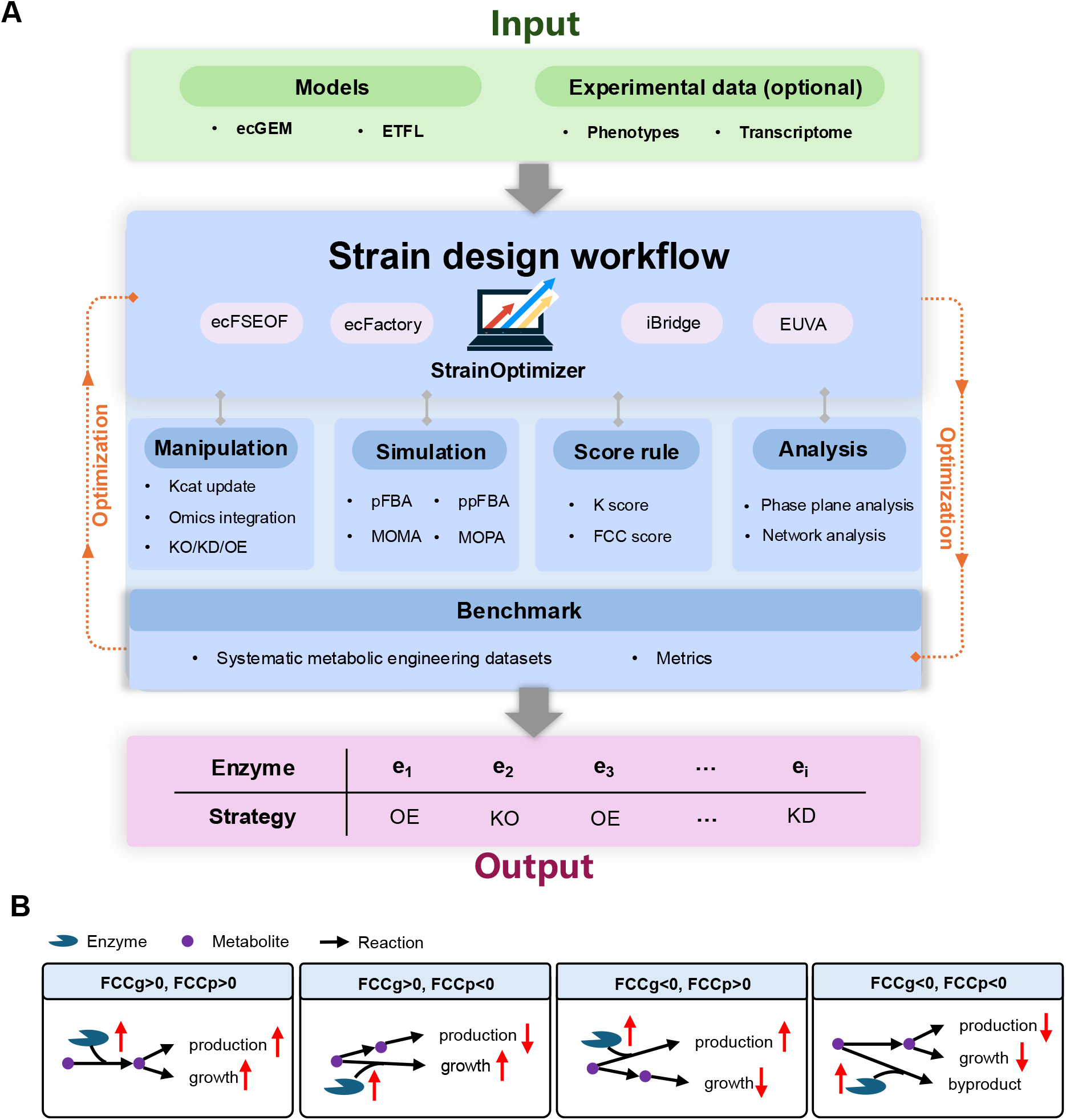
The computational framework of StrainOptimizer. A. Architecture of the modular StrainOptimizer platform. B. Schematic of the Flux Control Coefficient (FCC) analysis used for target identification. FCCg and FCCp represent the flux control coefficients for cellular growth and product synthesis, respectively.

Our previously developed ecFactory algorithm demonstrated integrative and multi-steps strain design method’s advantage in target prediction by integrating enzyme-constrained FSEOF(ecFSEOF) with enzyme abundance variability analysis to ensure predictive robustness and minimizing the number of false positives among predictions (*17*). Here, we extended its capabilities in two significant ways. First, building on the principle that metabolic objectives are affected by the abundance of key enzymes, we incorporated analysis of flux control coefficients (FCC) (*27*) as an optional scoring and screening method for targets (Figure 1B). Second, while the original ecFactory was limited to ecGEMs, we adapted the algorithm to be compatible with the more mechanistically complex ETFL models.

To enable rigorous algorithm assessment, we established a high-quality benchmark dataset. Recognizing that the effects of genetic modifications can vary significantly even within the same species due to the differences in the host strain’s genetic background (*28*–*32*), we curated our dataset with high stringency. For each of the five selected systematic metabolic engineering case studies in *Saccharomyces cerevisiae*, experimentally validated targets were sourced from a single paper to ensure a consistent genetic context. This included targets for the production of 2-phenylethanol (*33*), spermidine (*34*), free fatty acids (*35*), sclareol (*36*), and heme (*15*). These products were selected for their diversity in precursor metabolites and a wide range of protein synthesis costs. In total, the benchmark dataset contains 111 experimentally validated genetic targets, with each case study encompassing 11-38 genetic targets.

Alongside a high-quality dataset, a comprehensive suite of evaluation metrics is equally important. To assess predictive accuracy, we employed two standard metrics: precision, which represents the proportion of true positives among all predicted targets, and experimental consistency (recall), which represents the fraction of all known positive targets that were identified. Beyond accuracy, a critical need for metabolic engineers is the discovery of novel targets at the genome-scale for further production improvement. We hypothesized that genetic targets topologically distant from the primary product biosynthesis pathway or with higher network connectivity are less obvious and potentially more impactful (*6*). Therefore, to quantify the novelty of predictions, we introduced two additional metrics: distance, shortest metabolic path length (minimum reaction steps) from a target reaction to either the substrate or the product, and connectivity, defined as the number of adjacent reactions linked to a target reaction. These metrics are designed to evaluate how insightful or non-obvious the predicted targets are (Methods).

In summary, the strainOptimizer platform, featuring enhanced algorithms coupled with our high-quality benchmark and comprehensive evaluation metrics, establishes a robust and integrated framework. This provides a crucial foundation for systematic optimization of various cell factories.

### Systematic evaluation of strainOptimizer in gene target prediction and filtration

Using the strainOptimizer framework, we systematically compared the performance of ecGEMs and EFL models for strain design and evaluated the efficacy of the new FCC scoring method for targets filtration. To ensure a direct and fair comparison with ecGEMs, we utilized the EFL model (a variant of ETFL that excludes thermodynamic constraints) (*13*), as it is less computationally intensive and more amenable to automated reconstruction for diverse industrial strains than full ETFL models.

Since ecFactory is an integrative, multi-step algorithm, we first analyzed the sets of targets generated at each stage: the initial prediction (L1), followed by two sequential filtering steps (L2 and L3), and our newly introduced FCC method (Figure S1). For both ecGEM and EFL models, we observed a clear trend: as the filtering level increases from L1 to L3, the average number of predicted targets decreases (L1: 69, L2: 50, L3: 16, FCC: 9) while average precision consistently improves (L1: 0.13, L2: 0.14, L3: 0.34, FCC: 0.50) (Figure 2A). This increase in precision comes at the cost of filtering out some true positives, leading to a decrease in recall (L1: 0.37, L2: 0.30, L3: 0.15, FCC: 0.14). Strikingly, our new FCC method achieved a higher precision than the most stringent L3 targets (L3 vs FCC: 0.34 vs 0.50) while maintaining a comparable level of recall (L3 vs FCC: 0.15 vs 0.13), primarily by identifying a smaller, more refined set of high-confidence targets. We then assessed the novelty of the predictions using our distance and connectivity metrics. We found that targets from the high-precision L3 and FCC methods had lower average scores for both connectivity (L1: 14, L2: 14, L3: 9, FCC: 10) and distance (L1: 6.6, L2: 6.7, L3: 6.7, FCC: 5.6), suggesting they represent more intuitive and easily discoverable genetic strategies. This creates a clear trade-off for the user (Figure 2B): for researchers seeking rapid initial improvements, the high-precision targets from FCC or L3 are ideal. Conversely, for experts working with highly engineered strains, the broader L1 set serves as a richer source for discovering novel targets to further enhance production.

**Figure 2.**
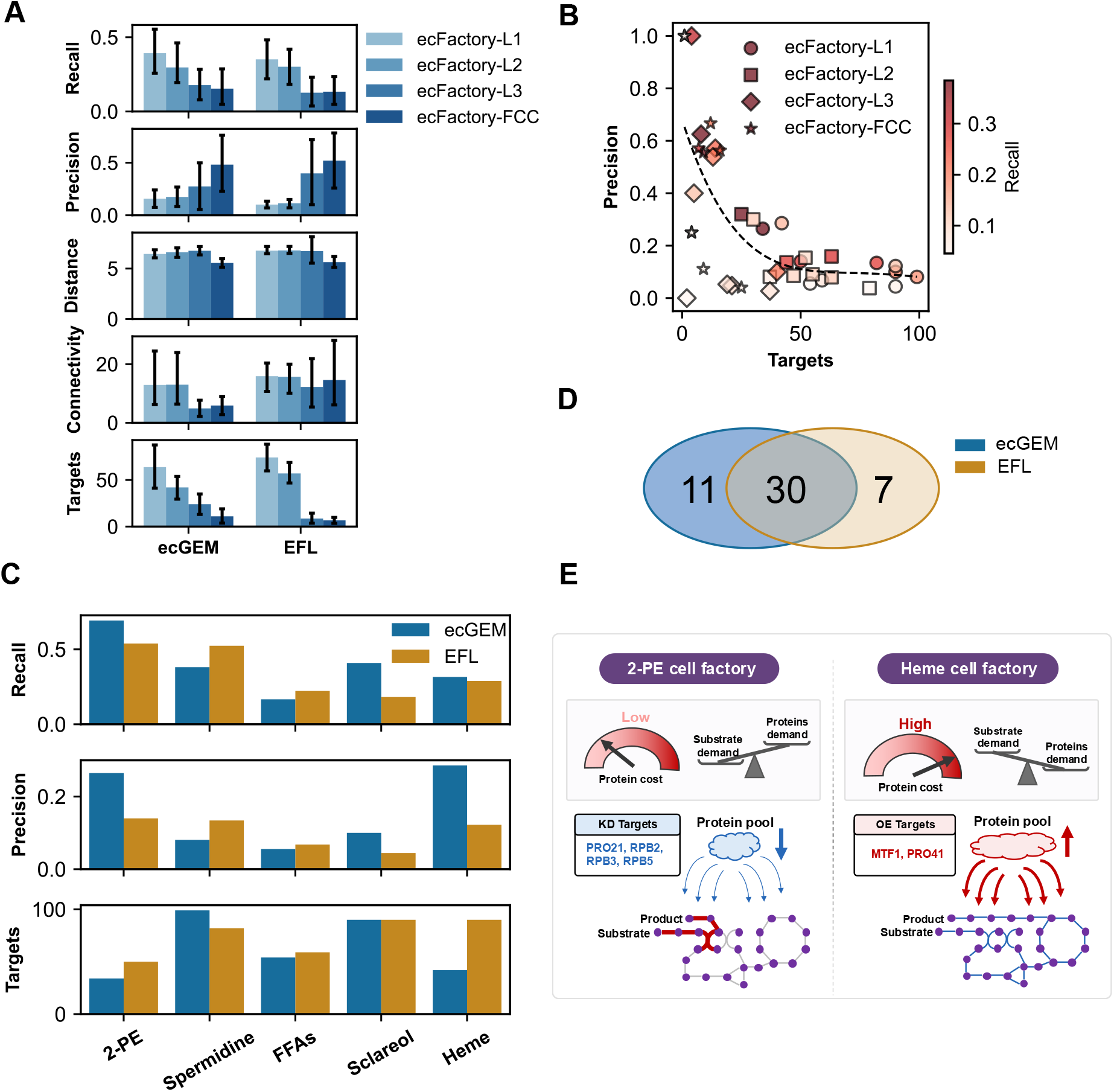
Performance comparison and complementary predictions of ecGEM and ETFL models. A. Performance metrics of the ecFactory algorithm at different filtering stages (L1, L2, L3, and FCC) for both ecGEM and EFL models. Error bars represent the standard deviation across five benchmark cases. B. The trade-off relationship between prediction precision and the number of predicted targets. Each point represents a prediction for one product at a specific filtering level. The color scale indicates recall. The dashed line represents a fourth-degree polynomial fit, illustrating the general trend that higher precision is achieved with smaller, more filtered target sets. C. Comparison of prediction performance between ecGEM and EFL models across the five different benchmark products. D. Comparison of aggregated ecFactory-L1 targets identified by ecGEM and ETFL models across all test cases. E. EFL model could capture systems-level targets for 2-PE and heme cell factory.

Next, we compared the overall performance of the ecGEM and EFL models. Our analysis revealed that the two models have distinct strengths for different products (Figure 2C, Figure S2A-C). For instance, the ecGEM demonstrated better performance for 2-phenylethanol (2-PE) (Recall/Precision: ecGEM: 0.69/0.26; EFL: 0.54/0.14), whereas the EFL model yielded better predictions for spermidine (Recall/Precision: ecGEM: 0.38/0.08; EFL: 0.52/0.13). Furthermore, the models often predicted different genetic targets, with each identifying unique targets that, when combined, resulted in a larger set of true positives (Figure 2D, Figure S2D-E). For example, in predicting targets for free fatty acids (FFAs), the ecGEM specifically identified upregulating the upstream pentose phosphate pathway (PPP) genes *ZWF1* and *GND1*. In contrast, the EFL model predicted that upregulating the downstream PPP enzyme *TAL1* was a better choice. The EFL model also successfully predicted that downregulating isocitrate dehydrogenase *IDH2* while upregulating the mitochondrial carrier *YHM2* would divert more carbon flux towards cytosolic lipid biosynthesis (Figure S3).

A key advantage of the EFL framework is its ability to identify targets beyond the metabolic network by accounting more cellular mechanisms like transcription and translation. The model predicted distinct, systems-level strategies based on the product’s specific resource demands (Figure 2E). For 2-PE, a product with a low protein cost (Table 1), the EFL model predicted that downregulating genes (*PRO21, PRB2, PRB3, PRB5*) involved in global transcriptional regulation was an effective strategy (*37*). This may reduce the overall cellular protein burden, thereby freeing up cellular energy and resources for precursor biosynthesis. In contrast, for heme—a product with a very high protein cost (Table 1, 2-PE vs heme: 0.2 vs 1.85 g_protein_/mmol_product_) that is primarily synthesized in the mitochondria—the model predicted a highly localized strategy: upregulating genes involved in mitochondrial gene expression system(*MRF1, PRO41*) (*37*). This finding highlights the model’s capacity to capture the crucial coupling between a biosynthetic pathway and the resource allocation of its primary subcellular compartment (*38*). Together, these results demonstrate the unique potential of ETFL and other multi-scale models to predict a broader range of non-intuitive targets for metabolic engineering.

**Table 1.** Benchmark dataset of representative systematic metabolic engineering cases.

| Product name | Targets number | Key precursor | Proteome cost<br>($\text{g}_{\text{protein}}/\text{mmol}_{\text{product}}$) | Ref. |
| --- | --- | --- | --- | --- |
| 2-phenylethanol | 11 | PEP/E4P | 0.02 | [33] |
| Spermidine | 22 | R5P/OAA | 0.12 | [34] |
| Free fatty acids | 18 | Ac-CoA | 0.20 | [35] |
| Sclareol | 22 | Ac-CoA | 0.25 | [36] |
| Heme | 38 | Suc-CoA/3PG | 1.85 | [15] |

### Subcellular proteomic constraints improve prediction precision

Cellular compartmentalization is a fundamental biological principle that allows cells to optimize resource utilization, mitigate biochemical incompatibilities, and enhance catalytic efficiency (*39*). It also has broad applications in metabolic engineering (*40*). Previous studies in systems biology have shown that integrating organelle-level constraints can improve the performance of cellular simulations (*12, 41*). Building on this, our group previously established a large-scale absolute proteome dataset for *Saccharomyces cerevisiae* including data from 30 different batch cultivation conditions (*10, 42, 43*). A key finding from this dataset was that major organelles consistently maintain a relatively stable mass fraction of the total cellular proteome(*44*). We therefore hypothesized that incorporating these experimentally observed ranges as compartment-specific protein pool constraints could enhance the predictive performance of strain design algorithms (Figure 3A).

**Figure 3.**
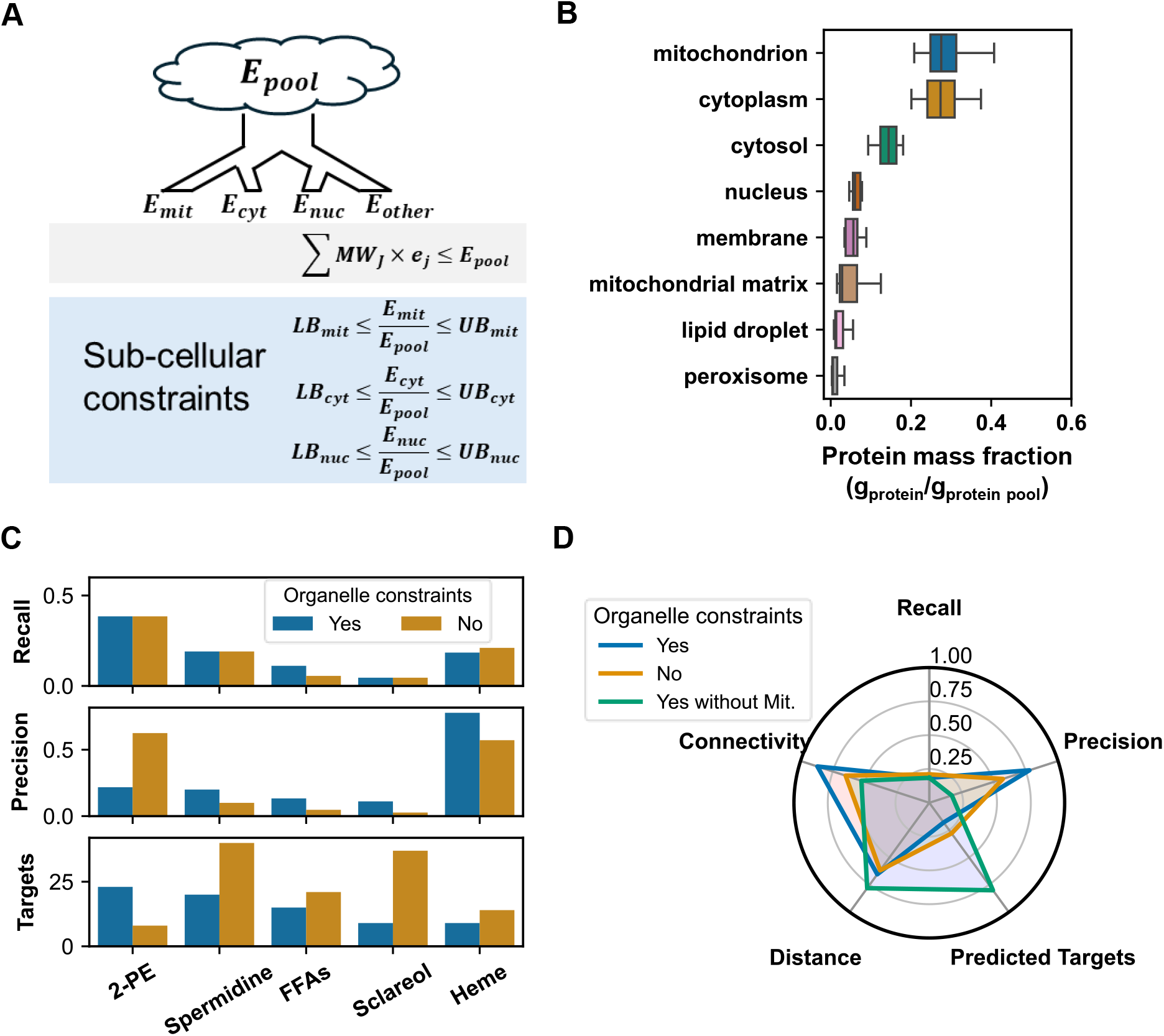
Subcellular proteomic constraints improve the precision of strain design predictions. A. The distribution of protein mass fractions for major organelles in *S. cerevisiae* across 30 different cultivation conditions. Protein mass fractions represent the ratio of a compartment’s total protein to the total cellular protein pool. B. Conceptual schematic of subcellular proteomic constraints in the models. C. Comparison of prediction performance with and without the application of organelle-level constraints across the five benchmark products. D. Ablation study for heme production, comparing key performance metrics under three conditions: with full subcellular constraints (Yes), with no constraints (No), and with all constraints except for the mitochondrial one (Yes without Mit.).

To derive constraints that are as relevant as possible to bioproduction conditions, we extracted a subset of this data from 30 batch cultivation conditions that most closely resemble fermentation environments to estimate the mass fraction ranges (Figure 3B). It is important to note, however, that these proteomics data was not collected under actual production conditions for the products. For the subsequent analyses, we selected the ecGEM as our base model due to its strong overall performance on our benchmark dataset and its lower computational demand compared to the EFL model.

Our results show that incorporating subcellular protein constraints significantly improves the prediction precision of the ecFactory algorithm (Figure 3C), though the effect was more moderate for the highly stringent FCC method (Figure S4A). Notably, this improvement was most evident for high-protein-cost products. For heme, the product with the highest protein cost, precision for the ecFactory algorithm increased from 0.57 to 0.78. Conversely, for 2-phenylethanol, the product with the lowest protein cost, the constraints offered no benefit (Figure 3C). These results suggest that the benefit of subcellular proteomic constraints may be more pronounced to the target product with high protein demand.

To further test this hypothesis, we performed a detailed case study on heme, which is primarily synthesized in mitochondria. We conducted an ablation analysis by removing only the mitochondrial protein constraint while retaining all others. The results show that precision dropped dramatically for both the ecFactory (from 0.78 to 0.18) and FCC (from 0.69 to 0.53) methods (Figure 3D, Figure S4B). This loss of precision was accompanied by a sharp increase in the number of predicted targets (from 9 to 40 for ecFactory, and from 13 to 17 for FCC), indicating that the mitochondrial constraint plays a key role in filtering false positives.

Analysis of the specific targets revealed that with the subcellular constraints applied, genes involved in cytosolic sterol biosynthesis (*ERG8, ERG12, ERG20, ERG13*) (*37*) were correctly excluded from the list of predictions for heme. This demonstrates that appropriate subcellular constraints can significantly improve the quality of predictions by enforcing biologically relevant resource allocation. However, their application requires careful consideration. For low-protein-cost products like 2-PE, the pathway’s resource demand may not be large enough to be meaningfully restricted by these constraints, rendering them ineffective in that context (Figure 3C). Furthermore, the use of inappropriate constraints, such as omitting a critical one as in our heme ablation study, can actively harm predictive accuracy (Figure 3D). As more high-quality, context-specific quantitative proteomics datasets for industrial microbes become available, the integration of organelle-level constraints into strain design pipelines offers significant promise for enhancing predictive performance.

### Protein-usage-based objective functions enhance predictive accuracy

The choice of objective function plays a crucial role in the model’s simulation of cellular phenotypes. Building upon the principles of Flux Balance Analysis (FBA)(*45*), two main classes of objective functions have emerged based on cellular resource allocation principles: those that minimize total resource usage(*46*) and those that minimize the adjustment of resources(*47*). While both have been widely applied (*11, 14, 17, 48*), they have not been systematically evaluated for strain design. The advent of enzyme-constrained models now allows these objective types to be extended from the flux level to the protein level, leading to pairs of analogous methods: minimizing total metabolic flux (pFBA) versus total protein usage (ppFBA), and minimizing metabolic flux adjustment (MOMA) versus protein usage adjustment (MOPA).

During metabolic reprogramming from growth to production states, cells undergo extensive resource reallocation. However, the dynamics of this reallocation differ significantly between metabolic fluxes and the proteome. To explore this, we simulated the shift from a maximal growth state to a maximal production state for each of the five products in our benchmark. The results consistently showed that the proteome is significantly more stable and less prone to drastic changes than the metabolic fluxome. Specifically, the distribution of protein usage changes exhibited a significantly smaller variance than that of metabolic fluxes changes(Figure 4A), indicating a lower overall demand for proteome reallocation. Furthermore, when considering extreme individual changes (log2 fold change > 5),far fewer enzymes showed extreme changes in protein usage compared to that of metabolic fluxes (8 vs. 32, respectively) (Figure 4B). This relative stability of proteome suggests that cells preferentially reroute metabolic fluxes (which are “cheap” to change) rather than executing costly, large-scale remodeling of their protein allocation. This implies that simulation objectives based on protein-level economy—such as minimizing total protein usage (ppFBA) or minimizing the adjustment of the proteome (MOPA)— are likely to be more biologically realistic and robust objectives for predictive modelling during *in silico* strain design.

**Figure 4.**
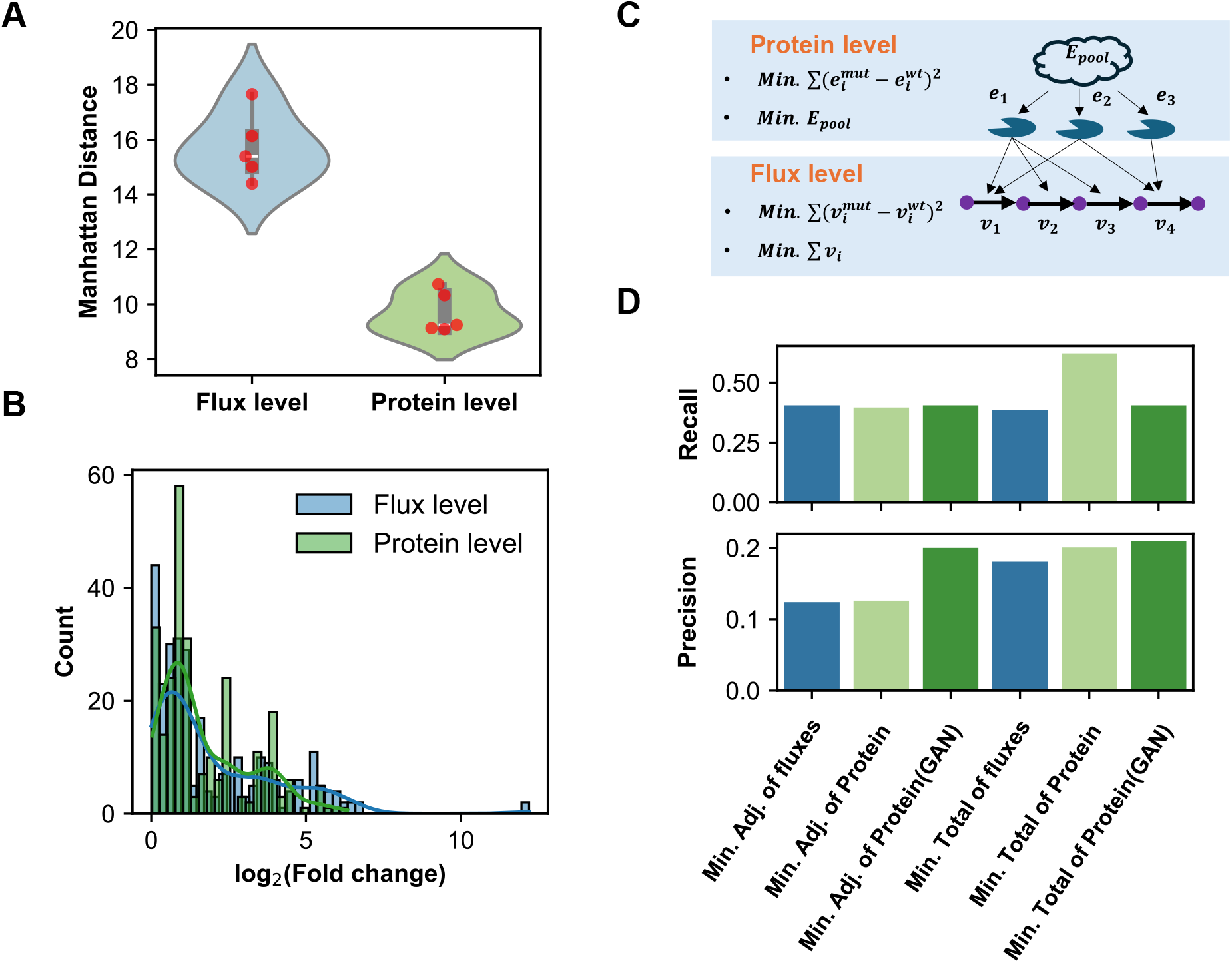
Protein-based simulation objectives improve the performance of strain design predictions. A. Comparison of Manhattan distances between growth and production states at flux and protein levels across five different products. B. Distribution of log2 fold changes in protein usage and metabolic flux when comparing growth versus production states across five product simulations. C. A schematic of the four simulation objectives evaluated, which are based on minimizing either total resource usage or resource adjustment at the protein and flux levels. D. Aggregated prediction performance metrics (recall and precision) for the four objective functions across all benchmark cases. Light and dark green shades distinguish predictions made using default versus improved k_*cat*_ parameter sets, respectively.

To systematically evaluate this, we implemented all four methods (pFBA, MOMA, ppFBA, MOPA) in our Simulation module (Figure 4C). The aggregated prediction results for our five case studies show that strategies based on optimizing protein allocation consistently achieved higher performance (Figure 4D). For the “minimize total resource” class, the protein-level objective (ppFBA) achieved significantly higher recall (0.62 vs. 0.39) and precision (0.20 vs. 0.18) than its flux-level counterpart (pFBA). For the “minimize adjustment” class, the difference was more subtle, with the protein-level objective (MOPA) showing only a slight improvement in precision over the flux-level objective (MOMA) (0.13 vs. 0.12). Overall, minimizing total protein usage (ppFBA) showed the best performance against the benchmark. While protein-level optimization showed a clear advantage, we hypothesized that its accuracy is related to the quality of the model’s kinetic parameters, particularly the genome-scale enzyme turnover rates (*k*_*cat*_). We repeated the analysis using an expanded set of *k*_*cat*_ parameters for *S. cerevisiae* that were curated using a generative adversarial network (GAN) (*49*). With these GAN-curated parameters, the predictive precision of the MOPA method improved significantly (from 0.13 to 0.20), becoming comparable to the ppFBA method (Figure 4D). However, for the ppFBA method, the optimized *k*_*cat*_ set paradoxically reduced its recall (from 0.62 to 0.41). A product-specific analysis revealed that while the updated parameters improved performance for low-protein-cost products (e.g., 2-PE and spermidine), they significantly hampered predictions for high protein-cost products like heme (Figure S5). This discrepancy likely arises from the context-specificity of the *k*_*cat*_ parameters. The GAN-curated set was optimized using proteome data from specific carbon- and nitrogen-limited cultivations (*49*). While *k*_*cat*_ values are often assumed to be constant, they can be highly condition-dependent. Consequently, applying these parameters to simulate high-burden pathways like heme synthesis—which operates under very different physiological demands—may impose unrealistic enzymatic constraints, leading to unreliable target predictions. This underscores that both the coverage and the contextual relevance of *k*_*cat*_ parameters are critical for *in silico* strain design.

In summary, protein-usage-based objective functions demonstrate a distinct advantage for strain design. Furthermore, the curation of model parameters like *k*_*cat*_ holds significant potential to improve predictions of gene targets for low-protein-cost products.

### StrainOptimizer could enable discovery of non-intuitive targets in an engineered overproducing sclareol strain

While our preceding analyses established a series of strain design pipelines suitable for various scenarios, their validation was based on existing benchmarks. This could not demonstrate strainOptimizer’s ability to guide the discovery of new targets in a practical metabolic engineering project. To address this, we applied the computational platform to SCX42, a sclareol-overproducing *S. cerevisiae* strain that has already undergone extensive metabolic engineering (*36*). The goal was to identify novel, non-intuitive genetic targets capable of further improving performance in this already optimized base strain.

We established a comprehensive workflow to generate predictions by integrating strain-specific phenotypic and transcriptomic data (Figure 5A). Predictions were generated by using both ecGEM and EFL models. Considering the Crabtre effect of *S. cerevisiae*, simulations were performed for both the glucose-consumption and ethanol-consumption phases of batch fermentation (*36*). Since our objective was to find non-obvious targets, we focused on the broad set of initial predictions from the ecFactory-L1 level, which were then filtered using product-specific FCC scores. This pipeline yielded 72 potential genetic targets. A pathway-based analysis of these candidates revealed that the largest cluster belonged to lipid metabolism—a finding consistent with previous studies on terpene engineering (*50, 51*), which have highlighted the competition for precursors between lipid and terpene biosynthesis (Figure 5B).

**Figure 5.**
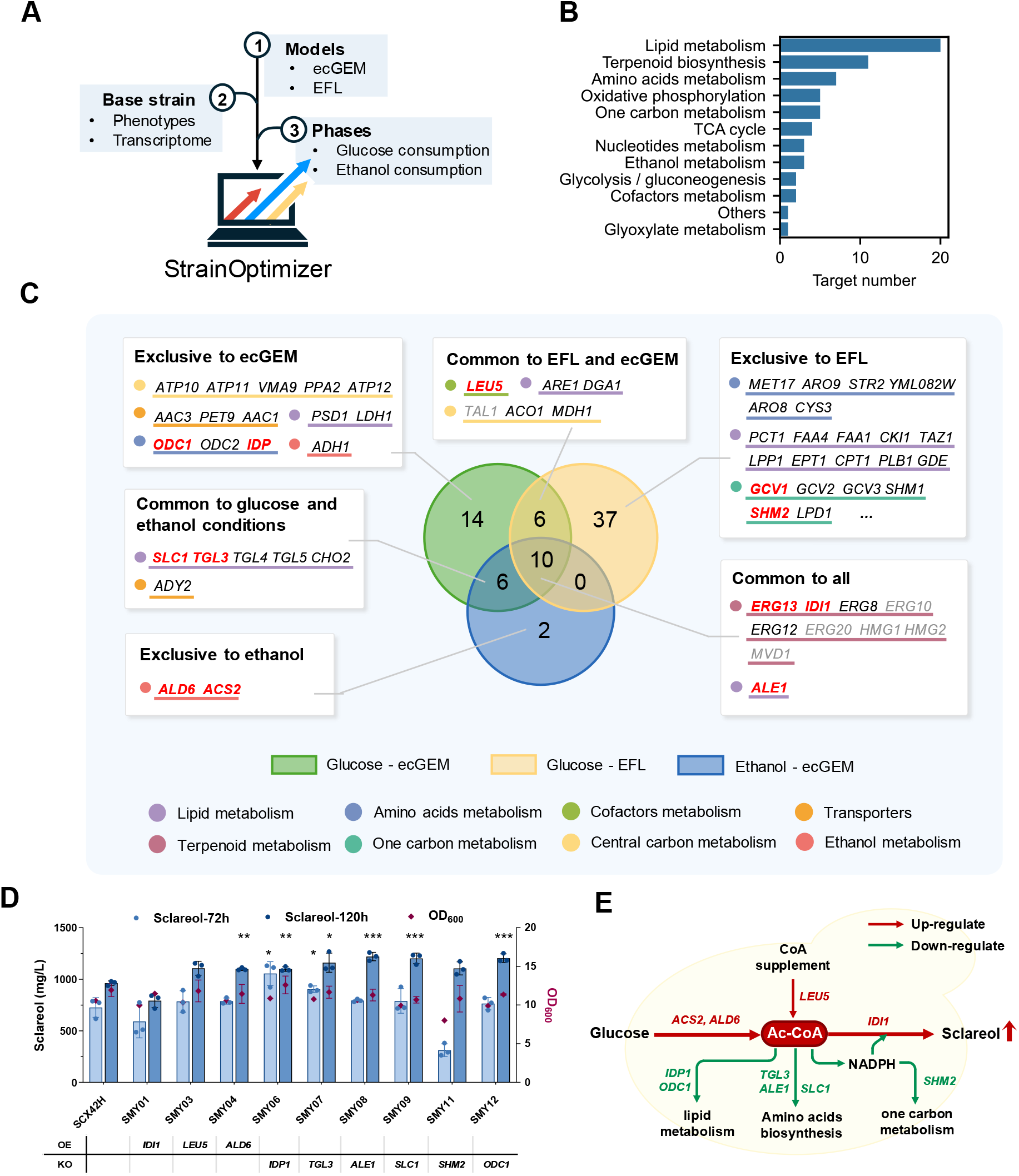
Application of strainOptimizer to identify novel targets for sclareol overproduction in S. cerevisiae. A. The computational workflow used to predict targets, which integrates strain-specific data (phenotypes, transcriptomics), metabolic models (ecGEM, EFL), and distinct fermentation phases (glucose and ethanol consumption). B. Pathway distribution of the potential genetic targets identified by strainOptimizer. C. Venn diagram categorizing the predicted targets based on the model and simulation condition in which they were identified. The background color of each section indicates the condition, while the color behind each gene name denotes its metabolic pathway. Genes selected for experimental validation are shown in red font; modifications pre-existing in the base strain are shown in blue. D. Experimental validation results for nine engineered strains. The bar chart displays sclareol titers at 72h and 120h, alongside the final cell density (OD_600_). Data are presented as mean ±SD from three biological replicates (n=3). OE denotes overexpression and KO denotes knockout. Error bars represent the standard deviation of biological triplicates. Asterisks indicate statistical significance compared to the control (statistical analysis was conducted by two-tailed t test, \**P*-value < 0.05, \*\**P*-value < 0.01, \*\**P*-value < 0.001). E. A simplified metabolic map illustrating the proposed mechanisms of the validated targets.

An analysis of the target origins revealed that the few targets common to all simulation conditions were concentrated in the terpenoid biosynthesis pathway itself (Figure 5C). These represent more intuitive targets, most of which have already been engineered in our base strain. Therefore, to identify novel candidates, we selected targets that were specific to individual models or simulation conditions. From this list, we chose 12 representative genes for experimental validation. Nine engineered strains were successfully constructed, and fermentation tests showed a 67% success rate, as six of the nine targets improved production. These effective targets increased the final titer by 14% to 26%, with the highest productivity enhanced by 45% (Figure 5D).

We first examined the fermentation profiles at 120 h, when the performance differences among engineered strains became clear. Six strains showed clear improvements, producing much higher levels of sclareol than the control strain. A closer look revealed that all effective targets were linked by a shared metabolic theme: improving how the cell supplies and allocates acetyl-CoA (Figure 5E). Among them, five knockout targets worked by reducing competition for common precursors acetyl-CoA, redirecting carbon flow toward the MVA pathway and boosting sclareol production. Specifically, knocking out gene *TGL3, ALE1*, and *SLC1* in lipid biosynthesis increased the final titer by 21-27%. Knocking out gene *IDP1* and *ODC1* in competing glutamate synthesis increased the titer by 14-25% (Figure 5D). In parallel, the overexpression of *ALD6* represented a different strategy, which strengthened the acetyl-CoA supply and led to a 14% increase in production. This finding suggests a general strategy that can be applied to improve the synthesis of other MVA-derived compounds.

To further evaluate performance beyond final titer, we assessed productivity at 72h, a phase of rapid sclareol accumulation. It is worth noting that the sclareol titer of the *idp1Δ*strain, which only had a 14% final titer increase, exhibited 45% increase in productivity during this earlier, rapid accumulation phase (Figure S6A, Figure 5D). This more pronounced effect on productivity suggests this target may hold even greater potential under industrial fed-batch conditions. Furthermore, to normalize for the impact of growth on performance evaluation, we calculated specific titers. This revealed that the *shm2Δ*, considered as a false positive target based on its comparable final titer, actually had a 27% higher specific titer (102.1 mg/L/OD_600_ versus 80.6 mg/L/OD_600_) than control strain SCX42H (Figure S6B, Figure 5D). This indicates the modification successfully enhanced per-cell production, but its severe negative impact on growth masked this benefit, suggesting a milder intervention like gene knockdown could be a successful strategy.

Overall, these findings illustrate how strainOptimizer can direct improvements in top-performing strains and guide the rational engineering optimization strategy.

## Discussion

In this study, we developed strainOptimizer, a comprehensive computational platform designed to systematically investigate and apply the principles of protein resource allocation for metabolic engineering. Our work addresses a critical bottleneck in the field: the need for accessible, robust tools that can translate the predictive power of advanced, enzyme-constrained models into actionable strain design strategies. By establishing a high-quality benchmark and a modular framework, we systematically evaluated how design performance is influenced by model type, subcellular constraints, and simulation objectives. We then demonstrated the platform’s practical utility an overproducing chassis strain, successfully identifying novel targets to improve sclareol production in an engineered yeast strain.

A central finding of our study is that the optimal application of protein resource allocation principles is context-dependent, requiring a multi-level approach. First, at the model level, our comparison between ecGEM (*11*) and EFL (*13*) models revealed their complementary strengths. For instance, in predicting targets for FFAs, ecGEMs were more effective at identifying targets in the pentose phosphate pathway, while the EFL model pinpointed key targets in the TCA cycle. Combining their predictions therefore yielded a more comprehensive and effective set of interventions (Figure S2). More importantly, EFL model could expand the target search space beyond metabolic enzymes. The prediction of downregulating global transcription for a low protein-cost product 2-PE versus upregulating mitochondrial expression for a high protein-cost, organelle-specific product heme showcases the unique capacity of multi-scale models to capture genetic targets related to systems-level resource trade-offs (Figure 2F). This suggests that a “one-size-fits-all” approach is insufficient; rather, an integrated strategy that leverages predictions from both model types is most practical for identifying a comprehensive set of potential genetic targets.

Second, at the constraint level, we demonstrated that adding biologically relevant constraints can significantly refine predictions. The incorporation of organelle-level proteomic constraints substantially improved prediction precision, particularly for heme, a product with high protein cost localized to the mitochondria (Figure 3C). Our ablation study, where removing the mitochondrial constraint reduced predictive accuracy, confirms that these constraints improve performance by filtering out a large number of biologically irrelevant false positives (Figure 3D). However, the failure to improve predictions for the low protein-cost, cytosolic product 2-PE underscores a critical principle: while additional organelle-level proteomic constraints improve the model’s precision in calculating metabolic flux, their practical benefits for refining strain design are realized only when they accurately reflect a biological resource limitation relevant to the specific product biosynthesis.

Finally, at the objective function level, our results clearly show the superiority of protein-based objectives over traditional flux-based ones. This is likely because proteome reallocation during metabolic shifts is more constrained and less variable than flux redistribution, making it a more tractable and robust basis for predictive modeling (Figure 4A, B). The ppFBA method, which minimizes total protein usage, consistently delivered the best performance. Furthermore, our finding that prediction accuracy can be further enhanced by using higher-quality kinetic parameters (*k*_*cat*_ values) highlights the deep interplay between model structure and parameterization. As these parameters become more accurate, the predictive power of protein-level optimization is poised to increase further. strainOptimizer represents a significant advancement over existing tools, including our previous work, ecFactory (*17*). It overcomes previous limitations by being re-engineered in Python for greater accessibility, supporting both ecGEM and ETFL models, incorporating new algorithmic modules like FCC analysis, and establishing a comprehensive benchmarking framework.

Furthermore, a key advantage of our work over recent algorithm-development studies (*6, 17*–*19, 52*) is the demonstration of practical utility beyond benchmark validation. Evaluating algorithms against existing modification data does not fully capture their ability to discover the novel, non-intuitive targets that experienced metabolic engineers require. To bridge this gap, we showcased a complete proof-of-concept workflow: from integrating strain-specific data and generating predictions to performing experimental validation. The successful application of strainOptimizer to an already highly engineered industrial sclareol-producing yeast strain (*36*) serves as powerful validation. The platform identified 6 of 9 novel targets that led to final titer by 14% to 26%, with the highest productivity enhanced by 45%, primarily by pinpointing critical metabolic nodes of competition for the acetyl-CoA precursor from lipid and amino acid biosynthesis.

Our study also highlights key limitations that point toward future research. The performance of advanced models and constraints is heavily dependent on the quality of available data, including proteomics and enzyme kinetic parameters. As more high-quality, condition-specific omics data become available, the predictive power of these models will improve (*25*). Future work should focus on expanding the benchmark dataset to more industrial host and more product, to assess the generalizability of our findings. Furthermore, strainOptimizer currently focuses on identifying individual genetic targets. Expanding its capabilities to predict synergistic combinatorial modifications is a critical next step for practical metabolic engineering (*53*–*55*). Finally, while the platform’s integrated, multi-step workflows enhance predictive power, they also introduce complexity. Integrating AI agent technology could simplify workflow design and lower the barrier to entry for non-expert users (*56*).

In conclusion, strainOptimizer provides a powerful, validated, and user-friendly platform that bridges the gap between the theoretical principles of cellular resource allocation and their practical application in metabolic engineering. By enabling a systematic, multi-level approach to strain design, it accelerates the development of high-performance microbial cell factories.

## Method

### Package Framework

The strainOptimizer platform is an open-source Python package engineered with a modular architecture. It is organized into five primary modules: Manipulation, Simulation, Analysis, Score Rule, and Strain Design Workflow. The Strain Design Workflow module serves as the core, orchestrating calls to the other four modules to predict genetic targets for a desired compound. The platform is compatible with multiple model types, including standard GEMs, ecGEMs, and ETFL models.

The Strain Design Workflow module currently includes four distinct algorithms: ecFSEOF, EUVA, ecFactory(*17*), and iBridge(*6*). These algorithms have been adapted from their original forms to incorporate advanced features, including the integration of transcriptomic data, the application of organelle-level proteomic constraints, and compatibility with multiple objective functions (pFBA, ppFBA, MOMA, and MOPA).

The Manipulation module provides functions for key model modifications, such as updating enzyme turnover rates (*k*_*cat*_), simulating gene knockout/knockdown/overexpression, and integrating multi-omics data. Transcriptomic data integration is implemented using the GIMME method (*57*) from the troppo package (https://github.com/BioSystemsUM/troppo).

The Simulation module includes four distinct objective functions for predicting cellular states, which are compatible with both ecGEM and ETFL models: flux-based optimization method: Parsimonious FBA (pFBA) (Minimizes the sum of all metabolic fluxes), MOMA(Minimizes the adjustment in protein usage according to reference state), and protein-based optimization method: protein-constrained pFBA(ppFBA) (Minimizes the total enzyme usage required to sustain a phenotype), MOPA(Minimizes the adjustment in protein usage according to reference state).

The score rule module provides two rules for scoring and ranking potential genetic targets. As described in our previous work(*17*), K-score is calculated for each gene based on the average flux score (ratio of mean flux in production vs. wild-type states) of all reactions associated with that gene. And flux control coefficients (FCC) are employed to quantify the control that each reaction exerts over cellular growth and production fluxes.

The Analysis module includes tools for evaluating and visualizing predictions. This includes phase plane analysis to visualize the production envelope of an engineered strain and network analysis functions to evaluate the metabolic distance and network connectivity of predicted target genes.

### Simulation Objectives

The first step of our strain design workflows needs to simulate cellular states under various production conditions for comparison against a wild-type reference. This process relies on different simulation objectives to estimate the distribution of metabolic fluxes and enzyme usage. The strainOptimizer platform supports four distinct objectives to compare flux-level and protein-level optimization strategies, all of which are compatible with both ecGEM and ETFL model frameworks.

pFBA is a two-step optimization that identifies the most flux-efficient phenotype. First, a standard FBA is performed to maximize a primary objective (*v*_*obj*_), typically the biomass production rate, to find its optimal value 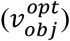:

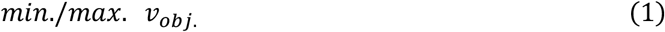

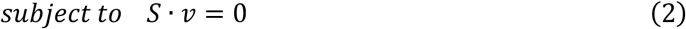

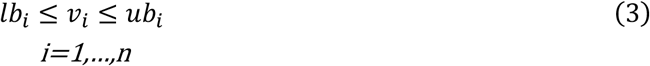

Here, S is the stoichiometric matrix, v is the vector of reaction fluxes, and lb and ub are the lower and upper bounds for each flux i.

In the second step, the primary objective is fixed as a constraint 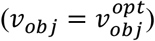, and the sum of absolute fluxes is minimized to find the most compact solution.

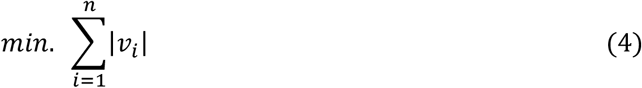

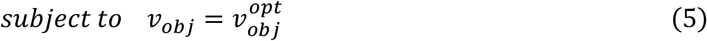

Analogous to pFBA, ppFBA finds the most protein-efficient solution. After the primary objective is maximized in the first step, the second step minimizes the total protein mass usage:

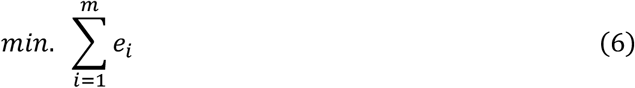

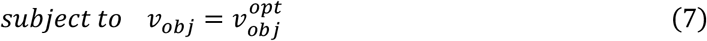

Where *e*_*i*_ represents the enzyme usage of reaction i and m is the total number of enzymes in the model. MOMA predicts the flux distribution of a perturbed strain by assuming it will make the smallest possible change from a reference (wild-type) flux distribution. It minimizes the sum of absolute differences in the metabolic fluxes between the reference and production conditions:

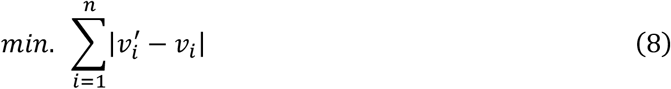

Where 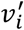 and *v*_*i*_ respectively represent the metabolic flux of reaction I under production and wild-type conditions.

MOPA applies the same principle as MOMA but at the proteome level. It requires reference enzyme usage distribution and minimizes the sum of absolute differences in the enzyme usage fluxes between the reference and production conditions.

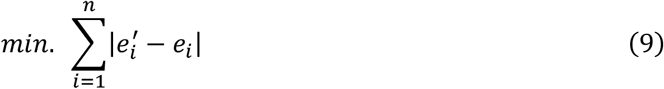

Where 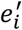and *e*_*i*_ respectively represent the enzyme usage of reaction I under production and wild-type conditions.

### Benchmark dataset collection

The benchmark dataset was curated from five high-quality, systematic metabolic engineering studies in *Saccharomyces cerevisiae*. For each of the five selected systematic metabolic engineering case studies in *Saccharomyces cerevisiae*, experimentally validated targets were sourced from a single paper to ensure a consistent genetic context. This included targets for the production of 2-phenylethanol (*33*), spermidine (*34*), free fatty acids (*35*), sclareol (*36*), and heme (*15*). These products were selected for their diversity in precursor metabolites and a wide range of protein synthesis costs. In total, the benchmark dataset contains 111 experimentally validated genetic targets, with each case study contributing between 11 and 38 genetic targets.

### Evaluation Metrics

To systematically assess the performance of the strain design algorithms, we used four distinct metrics. The first two evaluate predictive accuracy, while the latter two quantify the novelty and non-obviousness of the predicted targets.

Precision measures the fraction of predicted genetic targets that are actual positives (i.e., experimentally validated). A high precision score indicates that the algorithm has a low false-positive rate. It is calculated as:

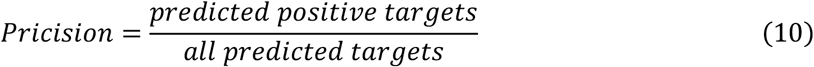

Recall measures the fraction of all known positive targets that the algorithm successfully identified. A high recall score indicates that the algorithm has a low false-negative rate. It is calculated as:

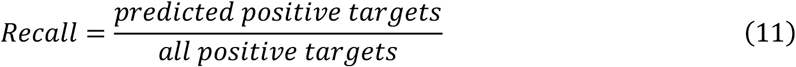

Distance is used to quantify the novelty of a predicted target. It is defined as the shortest metabolic path length (i.e., the minimum number of reaction steps) from a target enzyme to either the primary substrate uptake reaction or the final target production reaction. Targets with a greater distance are considered more non-intuitive, as they are further removed from the direct production route.

Connectivity is defined as the number of adjacent reactions linked to a target reaction. This metric is calculated from a reaction-metabolite graph that is first pruned to focus on the flow of the carbon backbone. The process is as follows:1. A graph is constructed where nodes represent either reactions or metabolites. 2. To isolate carbon-transferring pathways, common currency metabolites (e.g., H2O, O2, ATP, NAD, NADP, CoA) are removed from the graph. 3. Based on this pruned graph, connectivity for a target reaction is calculated as the sum of its upstream reactions (those that produce its substrates) and downstream reactions (those that consume its products).

### Flux Control Coefficient (FCC) analysis

To probe the control exerted by a specific enzyme on a given reaction flux, we conducted Flux Control Coefficient (FCC) analysis. The FCC of an enzyme i on a reaction flux v_j_ was calculated by simulating the product formation rate before and after a two-fold upregulation of the enzyme’s usage:

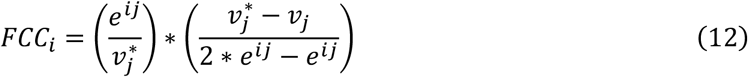

where v_j_ represents the original flux for reaction j in a reference flux distribution; e^ij^ is the enzyme usage for the enzyme i in reaction j; and 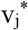 is the resulting flux for reaction j after perturbation. For FCC calculations with respect to production (FCCp), a value of FCCp>0 indicates a target for upregulation, while FCCp<0 suggests a target for attenuation.

Simulations were performed using minimal medium with glucose as the sole carbon source, with the glucose uptake upper bound constrained to 10 mmol/gDW/h. A two-round optimization approach was employed: First, the specific growth rate was fixed at 0.2 h^−1^ and product formation was constrained to 50% of the maximum product rate. Non-growth associated maintenance (NGAM) was maximized as the objective function using parsimonious flux balance analysis (ppFBA) optimization. Second, the target enzyme’s usage reaction lower bound was increased to twice its original value. Using the protein allocation profile from the first round as reference, MOPA was performed to obtain the secondary flux distribution and extract the 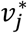.

### Calculation of Theoretical Protein Cost

The theoretical minimal protein cost for a product was defined as the total protein mass required to produce one unit of the product (g Protein / mmol product), calculated as described previously (*9*).

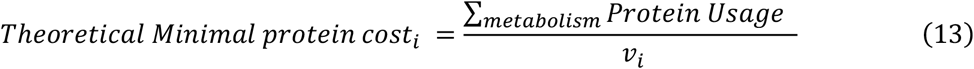

Where *v*_*i*_ is the flux of the objective, ∑_*metabolism*_ *Protein Usage* is the total proteome usage involved in metabolism. A two-stage optimization was employed: first, the product synthesis rate was maximized to find the theoretical maximum *v*_*i*_; second, this rate was fixed as a constraint, and the total protein pool was minimized.

### Implementation of Subcellular Proteomic Constraints

To incorporate organelle-level constraints, protein abundance data from various studies were first standardized to a common unit of mass fraction, referred from our recent work(*44*).

The total mass fraction for each organelle was calculated by summing the mass fractions of all proteins annotated to that compartment from the SGD database. The experimentally observed range (minimum and maximum) of these total mass fractions for each organelle across different conditions was then used to set the lower and upper bounds for the subcellular proteomic constraints in the model.

### Prediction of Novel Targets for Sclareol Production

1. Strain, Data Collection, and Model Preparation The base strain for this study was SCX42, a high-yielding sclareol-producing *S. cerevisiae* strain previously engineered in our laboratory. Strain-specific physiological data were collected during the exponential growth phase, including the specific rates of glucose consumption, sclareol production, and growth. Additionally, transcriptomic data were obtained from two key fermentation phases: the mid-exponential (glucose-consumption) phase and the ethanol-consumption phase (*36*). The ecGEM and EFL models were used. The heterologous sclareol biosynthesis pathway was manually curated and integrated into both models.
2. Model Constraining and Target Prediction The models were constrained differently for each fermentation phase to reflect the strain’s physiology. For the glucose-consumption phase, the specific substrate uptake rate was set to 11.5 mmol/gDW/h based on experimental data, and the measured specific growth rate was used to calculate the reference flux distribution. For the ethanol-consumption phase, where experimental data was unavailable, the substrate uptake rate was set to 1 mmol/gDW/h, and the reference growth rate was estimated as 49% of the maximum theoretical growth rate. For both phases, the corresponding transcriptomic data were integrated using the GIMME algorithm to prune the metabolic network and refine the solution space for reference condition calculation. Using these constrained models, *in silico* predictions were performed for both phases using the ecFactory algorithm with the ppFBA objective function. This process yielded four distinct sets of predictions (one for each combination of model and phase).
3. Target Integration, Filtering, and Selection for Validation The four sets of predictions were integrated and filtered according to the following criteria to generate a high-confidence list of candidates:
  1. Targets with conflicting modification directions (e.g., suggested for both overexpression and knockout) across the different sets were removed.
  2. The Flux Control Coefficient for production (FCCp) was calculated for each target. Overexpression (OE) targets with an FCCp < 0 and knockout/knockdown (KO/KD) targets with an FCCp > 0 were excluded.
  3. Essential genes, as defined by the model, were removed from the list of potential KO/KD targets.

This filtering process resulted in a final list of 72 targets. Due to limited experimental throughput, a subset was selected for validation. The selection was prioritized to identify novel targets outside of central metabolism and the primary terpenoid pathway, while also ensuring diversity across different metabolic pathways, originating models, and simulation phases. In total, 12 targets were chosen for experimental validation: *ERG13, IDI1, ALE1, ODC1, IDP, GCV3, SHM2, SLC1, TGL3, LEU5, ALD6*, and *ACS2*.

### Strain, media and cultivation

All strains developed for sclareol production in this study are detailed in Supplementary Table 1. *Escherichia coli* strain DH5α served as the host for all plasmid construction and amplification procedures, which were cultivated at 37 °C in Luria-Bertani medium (10 g/L tryptone, 5 g/L yeast extract, and 10 g/L NaCl). *S. cerevisiae* strains were routinely cultivated at 30 °C in YPD medium (10 g/L yeast extract, 20 g/L peptone, 20 g/L glucose) or SD medium (0.67% yeast nitrogen base without amino acids, 2% glucose with uracil (20 mg/L) if needed as previously described (*36*). The selection of transformants was conducted on SD medium solidified with 20 g/L agar. Plasmid curing was facilitated using YPD medium solidified with 20 g/L agar and supplemented with 5-fluoroorotic acid (5-FOA). For sclareol production, fermentations were carried out in a Delft minimal medium [38], which contained 2.5 g/L (NH4)2SO4, 14.4 g/L KH2PO4, 0.5 g/L MgSO4·7H2O, 20 g/L glucose. It was further supplemented with 2 mL/L of a trace metal solution, 1 mL/L of a vitamin solution and 60 mg/L uracil.

Inoculum preparation for shake-flask sclareol production involved growing engineered strains in Delft minimal medium for 18 hours. Preculture cells were harvested by centrifugation at 1200 × g, 5 min. The cell inoculated to fresh Delft minimal medium at a starting optical density of 0.2 at 600 nm. Cultures grew for 120 h at 30°C under 220 r.p.m. Samples enabled measurement of final biomass and sclareol levels.

### Genetic Manipulation

Genome editing in *S. cerevisiae* was carried out using a CRISPR/Cas9 system. The process included guide RNA (gRNA) design, assembly of gRNA expression vectors and donor fragments, transformation of yeast cells, and genotype confirmation.

Information on all gRNA-expressing plasmids and primers is provided in Supplementary Tables 2 and 3. gRNAs were designed to target the initial 5’ region of the open reading frame (ORF) of each target gene. The donor DNA for gene overexpression and seamless gene knockout was constructed by overlap extension PCR (OE-PCR). For scarless deletions, 300 bp sequences upstream and downstream of the target locus were amplified and joined directly. For targeted integration, cassettes contained promoter, gene, and terminator were inserted between these homologous regions. Yeast transformations were performed using the lithium acetate protocol. The gRNA plasmid and donor DNA (500 ng each) were co-transformed into the parental strain SCX42H. Transformants were selected on SD agar and incubated for 3 days at 30°C. Correct transformants were confirmed by colony PCR and verified clones were subsequently cultured on 5-FOA plates to remove the editing plasmid.

### Product Extraction and Quantification

Sclareol extraction and quantification followed a prior method [38] involving GC-MS analysis after liquid-liquid extraction with hexane that included 40 mg/L sclareolide as internal standard. Minor adjustments were applied. Pipette tips underwent trimming before sample withdrawal in cultures showing solid precipitates to promote accurate sampling. Extraction dilution 20-fold using hexane with 40 mg/L sclareolide. Analysis occurred on a ThermoFisher Scientific GC-MS setup featuring a DSQII mass spectrometer and Zebron ZB-5MS GUARDIAN capillary column, and the established program guided the process [38]. Xcalibur software handled data collection and quantification.

## Code and data availability

All codes and data are accessible at https://github.com/hongzhonglu/strainOptimizer.

## Acknowledgements

This work is supported by Shanghai Municipal Science and Technology Major Project, grant 2022YFA0913000 from the National Key Research and Development Program of China, and grants 22208211, 22378263, and 22425807 from the National Natural Science Foundation of China.

## Author contributions

H.L. and Y.J.Z. designed the research. H.W. performed algorithm development, data analysis and draft the original manuscript. M.Z. performed the experimental validation, analyzed the data. C.Z., S.H., W.L., R.Z. assisted in algorithm development. H.L., Y.J.Z., H.W., M.Z. revised the manuscript. All authors reviewed and approved the manuscript. H.L and Y.J.Z. obtained the funding for this study.

---

**Figure S1.**
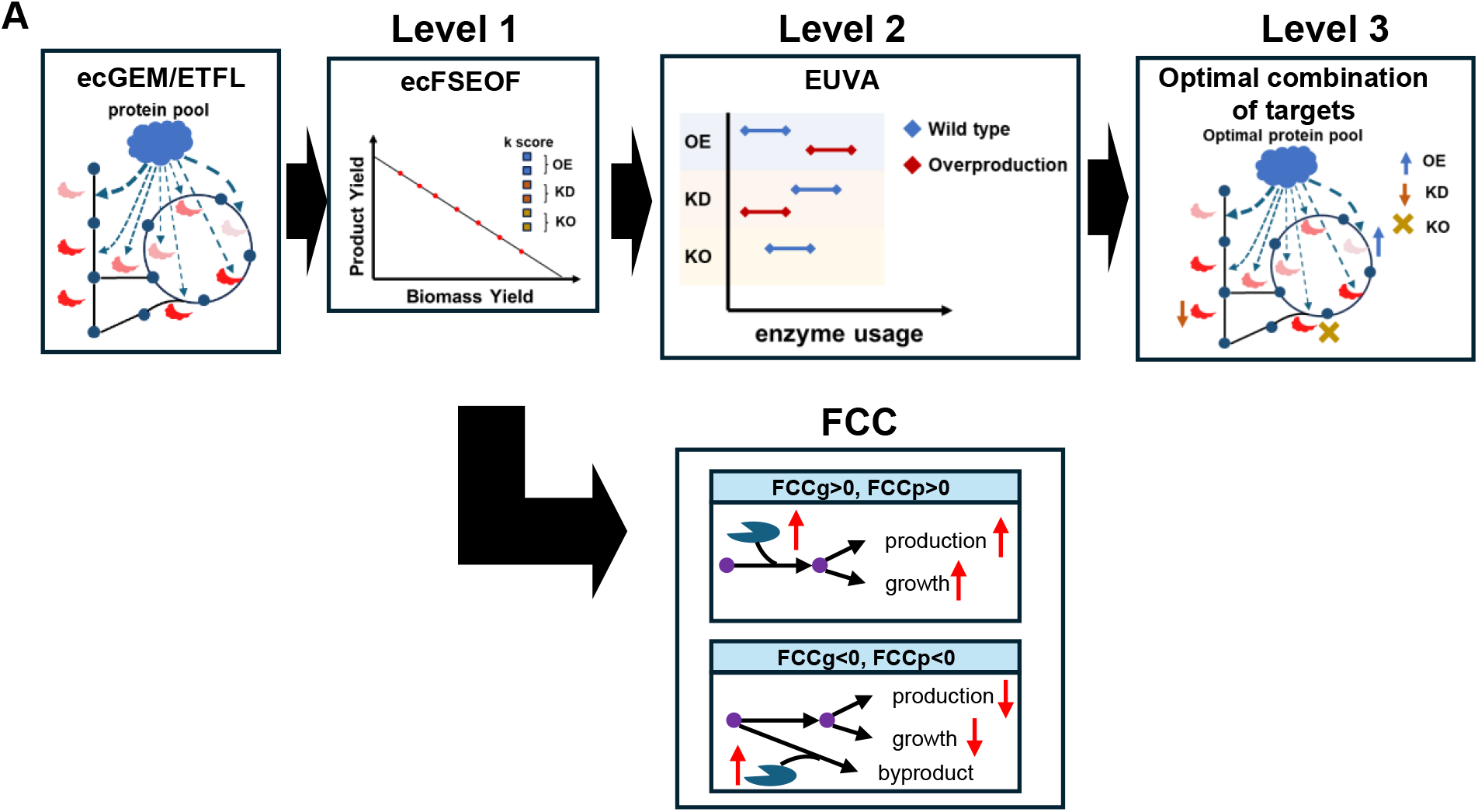
Schematic diagram of the enhanced ecFactory strain design pipeline.

**Figure S2.**
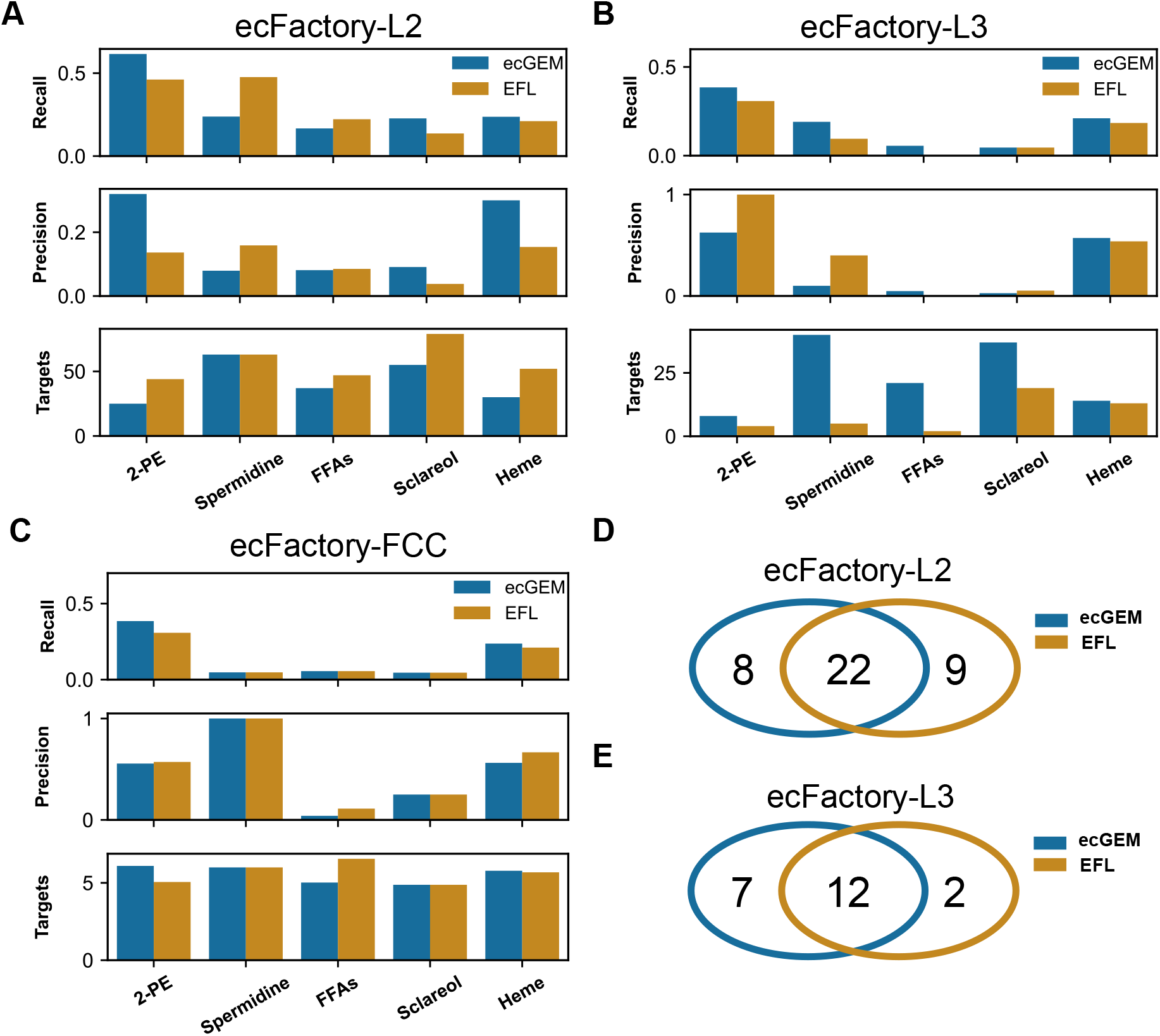
Performance comparison and complementary predictions of ecGEM and ETFL models. Comparison of prediction performance between ecGEM and EFL models across the five different benchmark products at the ecFactory-L2 level (A), ecFactory-L3 level (B), ecFactory-FCC level (C). Comparison of aggregated true positive targets identified by ecGEM and ETFL models across all test cases at ecFactory-L2 level (D) and ecFactory-L3 level (E).

**Figure S3.**
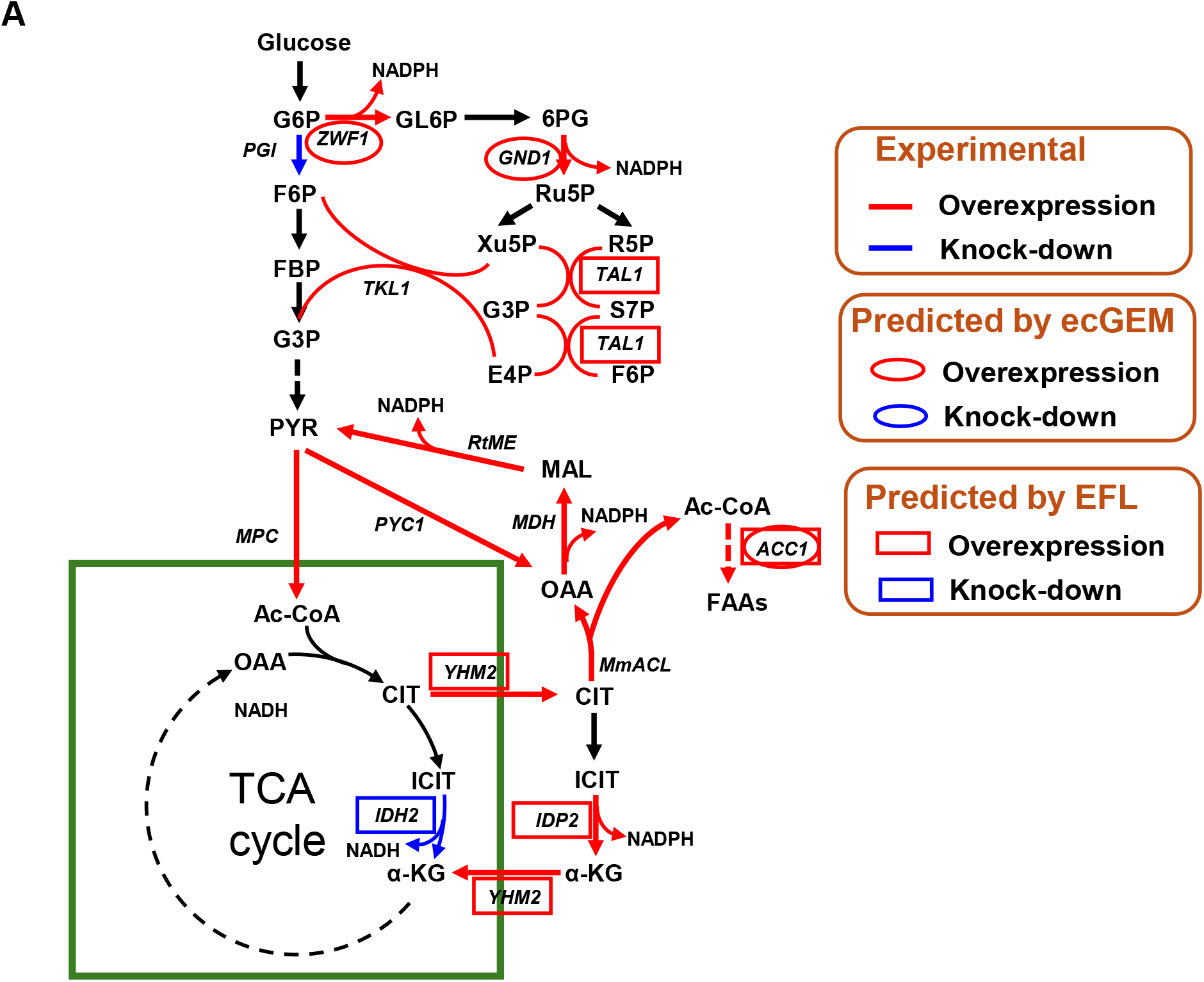
Complementary genetic targets for free fatty acid (FFA) production predicted by ecGEM and EFL models.

**Figure S4.**
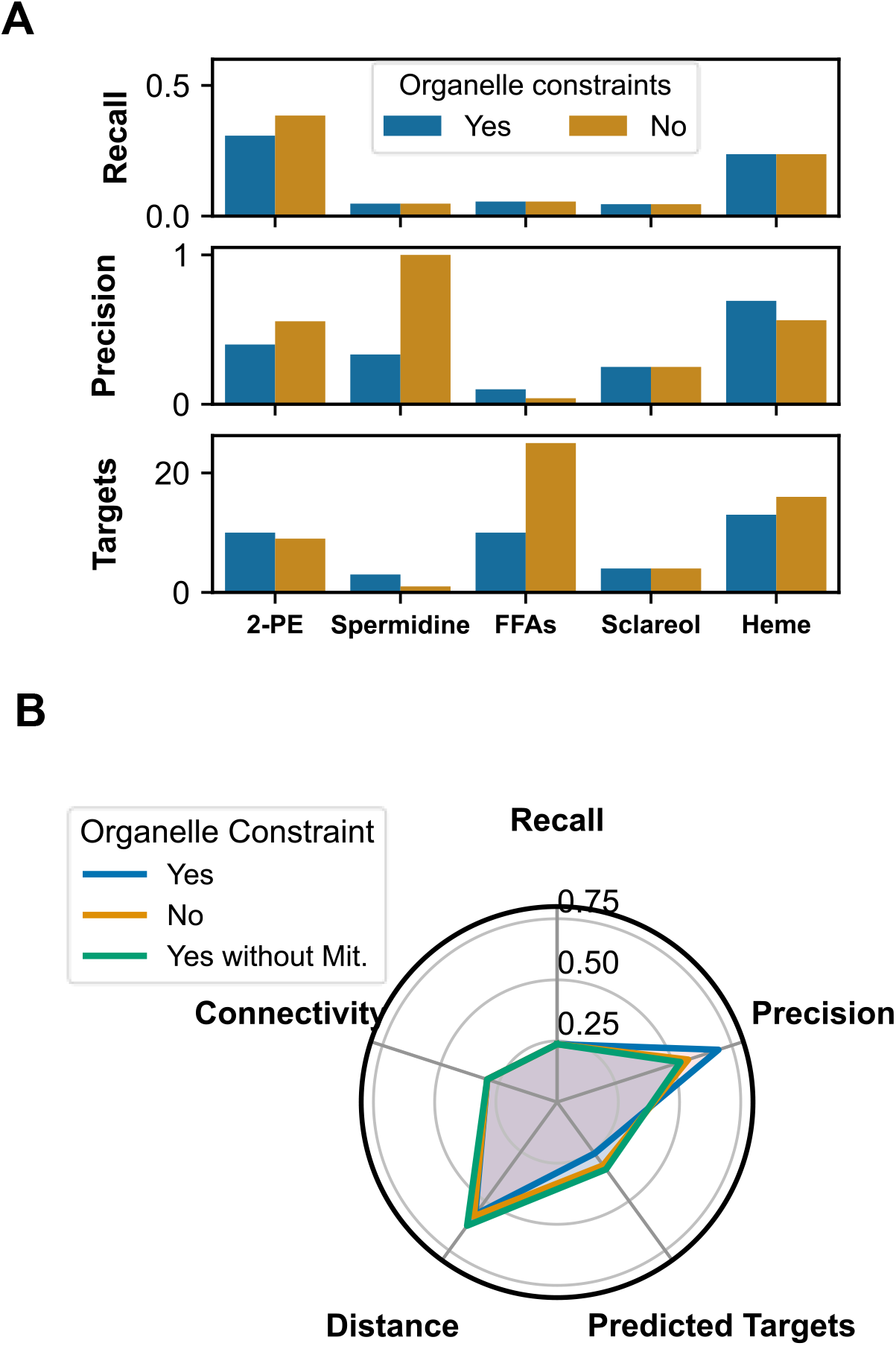
Effect of subcellular proteomic constraints on the FCC prediction method. A. Comparison of prediction performance with and without the application of organelle-level constraints across the five benchmark products for the FCC method. B. Ablation study for heme targets prediction by the FCC method, comparing key performance metrics under three conditions: with full subcellular constraints (Yes), with no constraints (No), and with all constraints except for the mitochondrial one (Yes without Mit.).

**Figure S5.**
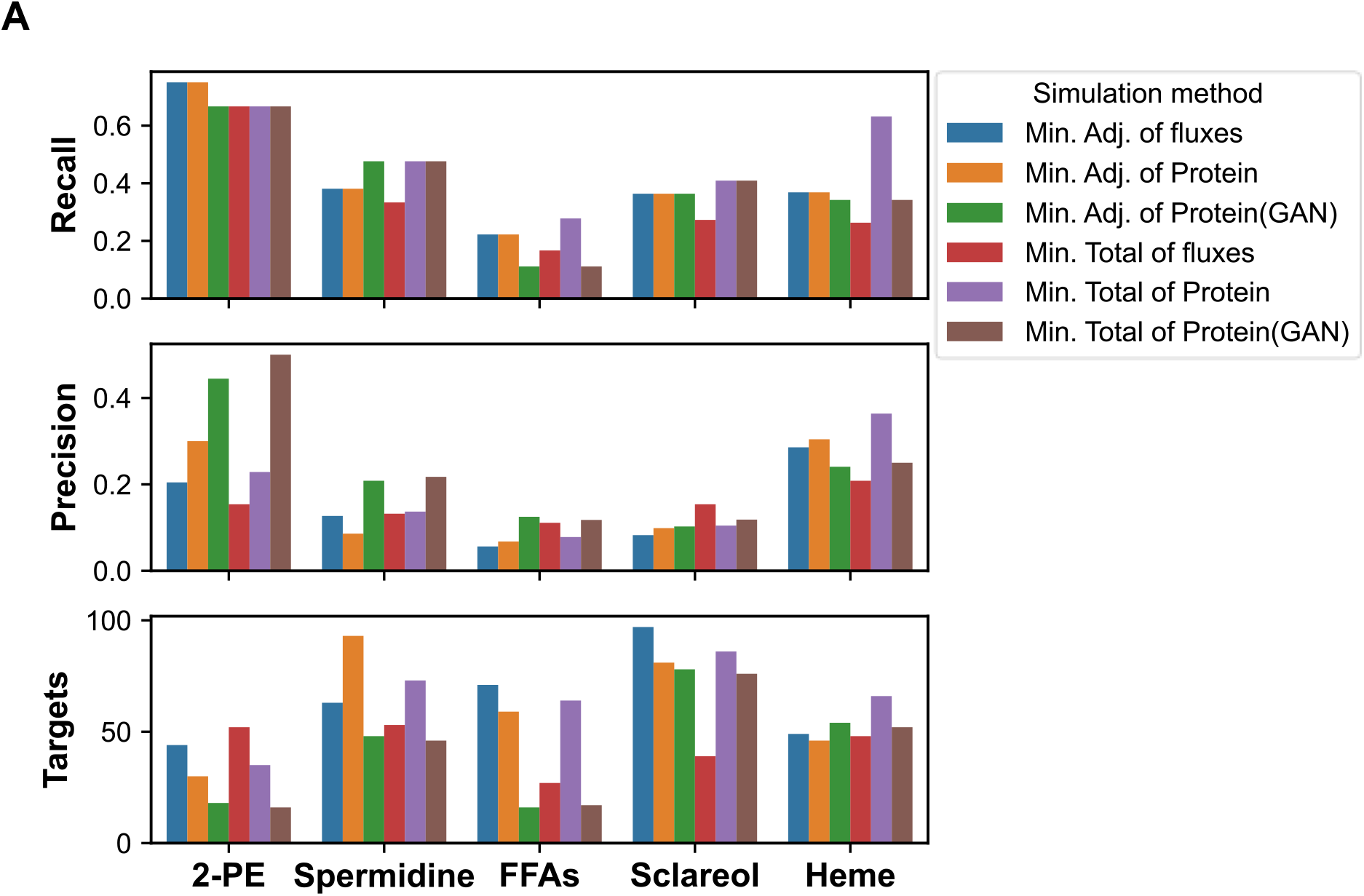
Product-specific performance comparison of simulation objective functions.

**Figure S6.**
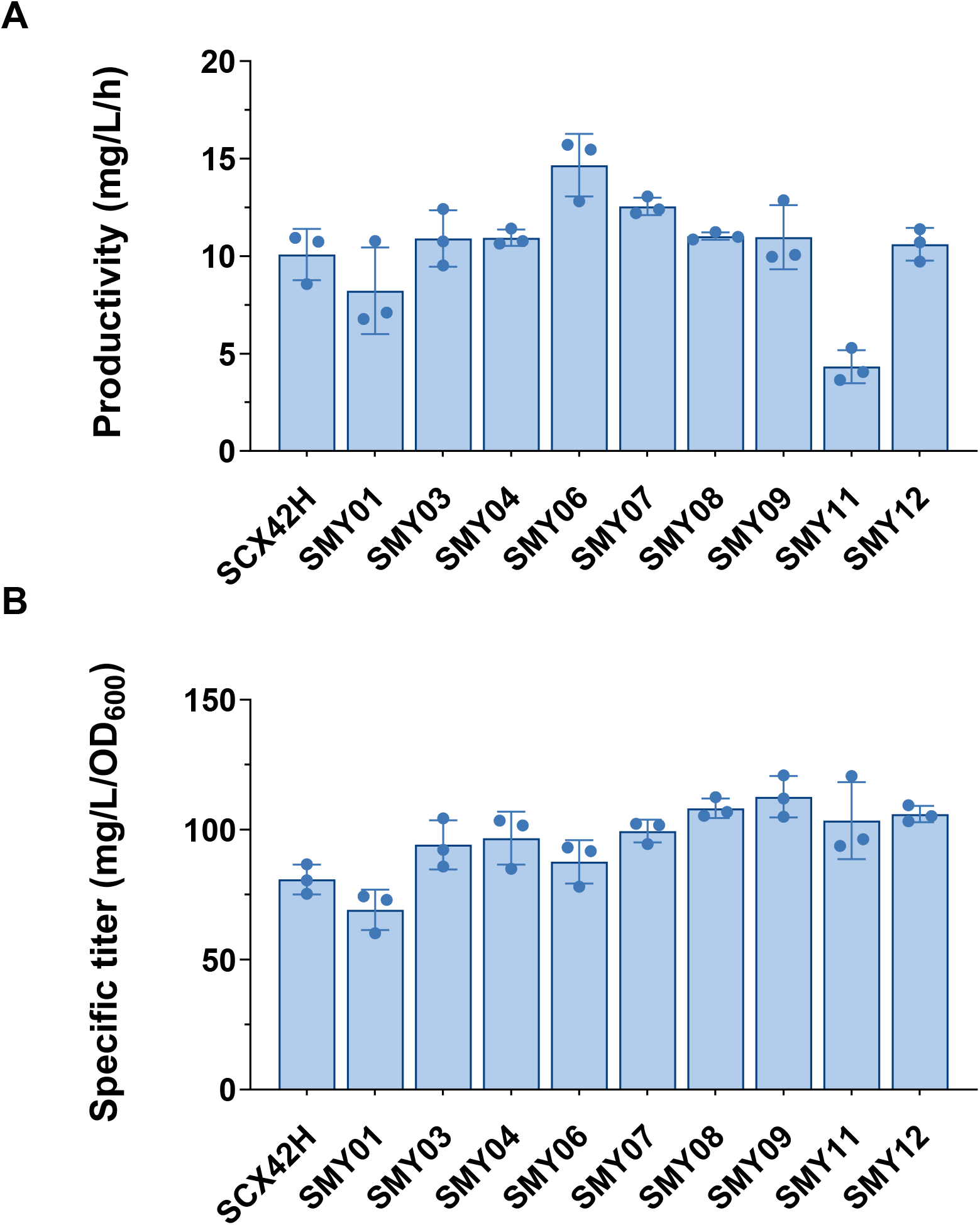
Experimental validation results for nine engineered strains. A. Sclareol productivity (mg/L/h) calculated from 0 to 72 hours. B. Final specific sclareol titer at 120 hours. Data are presented as mean ±SD from three biological replicates (n=3).

